# Senescent cells impair fracture repair through elevating ubiquitin-proteasome system activity in aged mice

**DOI:** 10.1101/2023.11.01.565138

**Authors:** Jun Zhang, Jiongnan Xu, Jiatong Liu, Brea Lipe, Tao Wu, Brendan F. Boyce, Jie Shen, Lianping Xing, Hengwei Zhang

## Abstract

Senescent cells accumulate in multiple tissues with aging. Depletion of senescent cells benefits the aging related disease, such as aging bone fracture. However, the molecular mechanisms by which senescent cells regulate their neighboring bone cells are still not well-known. We reported that proteasome inhibitor enhanced fracture repair in aged mice. Senescent cells are major source of chronic inflammatory cytokines, which in turn induced protein ubiquitination. We reported that PDGFRβ was one of the highly ubiquitinated proteins in mesenchymal progenitors (MPCs) and TGFβ was the most increased SASP. In the current study, we found TGFβ induced PDGFRβ ubiquitination and proteasomal degradation through its E3 ligases. TGFβ neutralizing antibody blocked the inhibited callus derived MPC growth and increased Ub-PDGFRβ by senescent cells, which could be further prevented PDGFRβ inhibitor. These findings suggested senescent cells derived TGFβ impaired fracture repair in aged mice through elevating ubiquitination of PDGFRβ. The discovery of TGFβ-PDGFRβ pathway triggered by senescent cells opens avenues for optimizing treatment strategies for aging related disease by combination with the ligand of PDGFRβ.

## Introduction

Cellular senescence was firstly reported by Dr. Hayflick and his colleagues as a permanent proliferative arrest^1^ and senescent cells were characterized by positive staining for senescent associated-b-gal (SA-b-gal) (pH=6) and expressions of altered gene expression patterns leading to a senescence-associated secretory phenotype (SASP)^2,3^. Senescent cells exhibit both beneficial and detrimental functions caused by different expression pattens of SASPs or their physical locations^4,5^. We^6^ and others^7,8^ have reported that increased senescent cells in bone inhibit the osteogenesis and promote osteoclastogenesis leading to osteoporosis and impaired fracture repair in aged mice. Depletion of senescent cells either by genetic or pharmaceutical approaches rejuvenated the aging related bone loss, arthritis and bone fracture^6-8^.

Although numerous molecular mechanisms involve in the inhibition of osteogenesis in aged mice, few studies reveal the targeting molecular mechanisms by which senescent cells contribute to the impaired osteogenesis. Dr. Farr’s group^7^ reported the decreased expression of sclerostin (Sost) after the depletion of senescent cells, which infer that senescent cells may inhibit the osteogenesis by inhibiting Wnt signaling through elevating the expression of Wnt signaling antagonist, Sost. Josephson’s group also reported that increased cellular senescence in mesenchymal progenitor cells decreased their stem/progenitor cell pool^9^. Therefore, there is a merit need to elucidate the molecular mechanism by which senescent cells affect their neighboring cells.

Ubuiquin-proteasome system (UPS) is recognized as the major pathway for regulated degradation of intracellular protein in proteasome^10^. Numerous studies reported that aged subjects have increased UPS activity^11,12^. We reported that aged mice have increased proteasomal degradation of Runx2 and JunB^12^. Inhibition of proteasomal protein degradation with proteasome inhibitor, Bortezomib enhanced fracture repair in aged mice^13^. Cellular senescence is closely associated with chronic inflammation under aging condition^9^. Inflammatory cytokines are key triggers for increased protein ubiquitination and degradation^14,15^. Therefore, we hypothesize that senescent cells impair fracture repair in aged mice through elevating UPS activity via SASPs.

## Results

### Senescent cells increase UPS activity

Increased senescent cells^6,7^ and UPS activity^11,12^ in aged mice have been reported. We also reported that both depletion of senescent cells by senolytic drugs and treatment of proteasome inhibitor enhanced bone fracture repair in aged mice^6^. To investigate the relationship between senescent cells and UPS, we examined the total Ub-protein in callus-derived mesenchymal progenitor cells (CaMPCs) treated with callus condition medium (CM) from vehicle or senolytic drug, dasatinib (D) + quercetin (Q) treated young and aged mice. Generations of CaMPCs and callus CM were standardized, and the effects of callus CM from aged mice (aged CM) are tested due to the existence of senescent cells in callus in our previous publication^6^. Western blots data showed that aged CM induced much more ub-proteins compared to young CM, which was blocked by D+Q treatment in aged mice (Fig. 1A). To examine whether the increased ub-protein are going to proteasome or lysosome pathway for degradation. CaMPCs were co-treated with young/aged CM and proteasome inhibitor, bortezomib (Btz) or lysosome inhibitor, chloroquine (CQ). The decreased CaMPCs growth and increased Ub-protein induced by aged CM can be reversed by Btz, but not by CQ (Fig. 1B&C). Data suggested that UPS serves a downstream signaling mechanism by which SCs regulating MPCs function.

**Fig.1.**
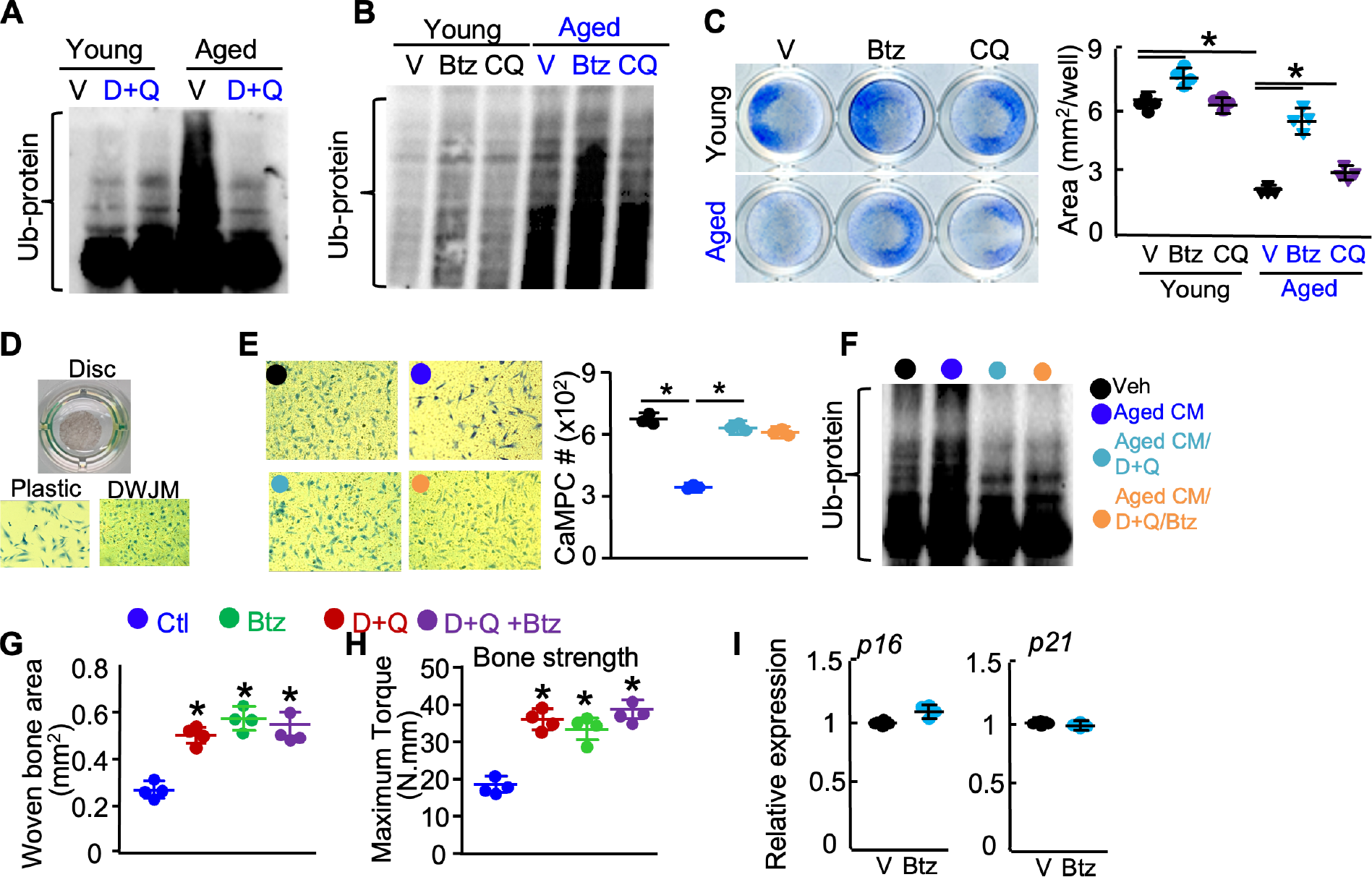
Aged CM increases UPS activity and reduces growth of CaMPCs, which is prevented by D+Q or Btz. CaMPCs were treated with young or aged CM ± D+Q or proteasome/lysosome inhibitor (Btz/CQ). Expression of total Ub-proteins by WB in D+Q-treated cells (A) and in Btz/CQ-treated cells (B). (C) CaMPCs were treated with young or aged CM ± Btz/CQ. Cell growth was assessed by methylene blue staining and positive cell area were measured by ImageJ. (n=4 wells). (D) Decellularized Wharton jelly matrix (DWJM) discs from the umbilical cord were used to 3D culture with CaMPCs in 96-well plate. After culture, cells on DWJM and plastic plate were fixed with 10% formalin and stained with methylene blue. (E-F) 3D culture of CaMPCs with aged CM ± D+Q ± Btz. Cells on DWJM were counted after staining. (n=3 wells). Separate CaMPC culture in 10cm dish was treated as in E were examined the total Ub-protein (F). (G&H) Aged mice were treated with Btz and/or D+Q. Callus was examined for woven bone area by histology at 10 dpf or bone strength by biomechanical testing at 28 dpf. n=4 mice. (I). p16 and p21 expressions in callus of aged mice treated with Btz were examined by qPCR. Data represent mean ± SD. 1-way ANOVA/Tukey test. \**p*<0.05.

### UPS involves in the effect of senescent cells on fracture repair in aged mice

To further examine the involvement of UPS in effect of SCs on fracture repair, we first used a well-established 3D culture system^16^ to examine whether the effect of SCs on CaMPC growth relied on regulating the downstream of UPS activity in a 3D micro-environment. Briefly, CaMPCs were cultured on a decellularized Wharton jelly matrix from the umbilical cord (Fig. 1D) and treated with different CM and Btz. Similar to results from Fig. 1B&C, aged CM inhibited CaMPC growth and increased total Ub-protein, which prevented by D+Q treatment on fractured mice (Fig. 1E). However, aged CM + D+Q + Btz can not give more cells compared to aged CM + D+Q (Fig. 1E), because additional Btz treatment did not prevent more protein degradation after reducing protein ubiquination by D+Q treatment (Fig. 1F). Furthermore, we treated aged mice with fracture surgery with D+Q ± Btz as we published before^6^. Histology analysis of woven bone in callus at d10 and biomechanical testing at d28, the gold-standard method to examine the fracture repair^17^, showed that D+Q or Btz alone can increase the new bone and bone strength, but they did not have addictive effect on fracture repair (Fig. 1G&H). To examine whether the lack of addictive effect is caused by the inhibition of senescent cells by proteasome inhibitor, although it was reported that proteasome inhibitor adversely induced cellular senescence by reducing the expression of hTERT and stabilizing pro-senescence factor, p53 and p21^18^, we examined the expression of senescent marker genes, p16 and p21 and did not find significant change in callus after Btz treatment (Fig. 1I).

### TGFβ1 induces ubiquitination and proteasomal degradation of PDGFRβ

Aged mice have impaired fracture repair with decreased osteogenesis and angiogenesis^19^. Proteasome inhibitor, Btz enhanced fracture repair in aged mice by increasing the number of MPCs and blood vessels (another manuscript to Biorxiv). Proteasome inhibitor has multiple targets in cells. We reported that tyrosine kinase receptor, platelet-derived growth factor receptor β (PDGFRβ) was the major target of Btz in C3H10T1/2 MPCs (another manuscript to Biorxiv). Meanwhile, PDGFRβ signaling plays a positive role both in osteogenesis and angiogenesis (ref). PDGFRβ protein expression was dramatically reduced in callus of aged mice. Btz enhances fracture repair in aged mice through inhibiting PDGFRβ degradation. However, the molecular connection between senescent cells and UPS-PDGFRβ is still not clear. We reported that TGFβ was the most increased SASP in callus of aged mice. Blocking of TGFβ with neutralizing antibody enhanced fracture repair in aged mice^6^. To investigate the mechanism by which senescent cells regulate the UPS activity and PDGFRβ degradation, we screened expressions of 7 key SASPs, related with osteogenesis and angiogenesis. We found only three of them are increased, including *Tgfβ, tnfα* and *Il-1* at mRNA level (Fig. 2A), but TGFβ at protein level has the most difference in all bone tissues between young and aged mice (Fig. 2B). CaMPCs were treated with different concentrations of TNFα, IL-1. Both TGFβ and TNFα decreased the expression of PDGFRβ, but TNFα level and change in callus between young and aged mice are minimal (Fig. 2C), so we chose TGFβ as the major SASP for the following mechanism studies. CaMPCs treated with TGFβ have increased Ub-PDGFRβ and decreased total PDGFRβ with a time-dependent manner (Fig. 2D). To examine whether TGFβ induced PDGFRβ degradation through proteasome or lysosome pathway, we treated with CaMPCs with TGFβ ± Btz ± CQ and found that TGFβ induced PDGFRβ degradation majorly through proteasome pathway (Fig. 2E&F). The Cbl family (Cbl, Cbl-b and Cbl-c) comprises the dominant RING finger ubiquitin E3 ligases for tyrosine kinase receptor protein degradation^20^. Cbl and Cbl-b mediates PDGF-induced PDGFRβ degradation^21^. We examined the expression of *Cbl* and *Cbl-b* and found both are increased in callus of aged mice. Treating CaMPCs with TGFβ in vitro also increased their expressions. Data suggested that TGFβ induced PDGFRβ proteasomal degradation through E3 ligases may represent a potential molecular mechanism by which senescent cells regulate CaMPCs function through UPS.

**Fig.2.**
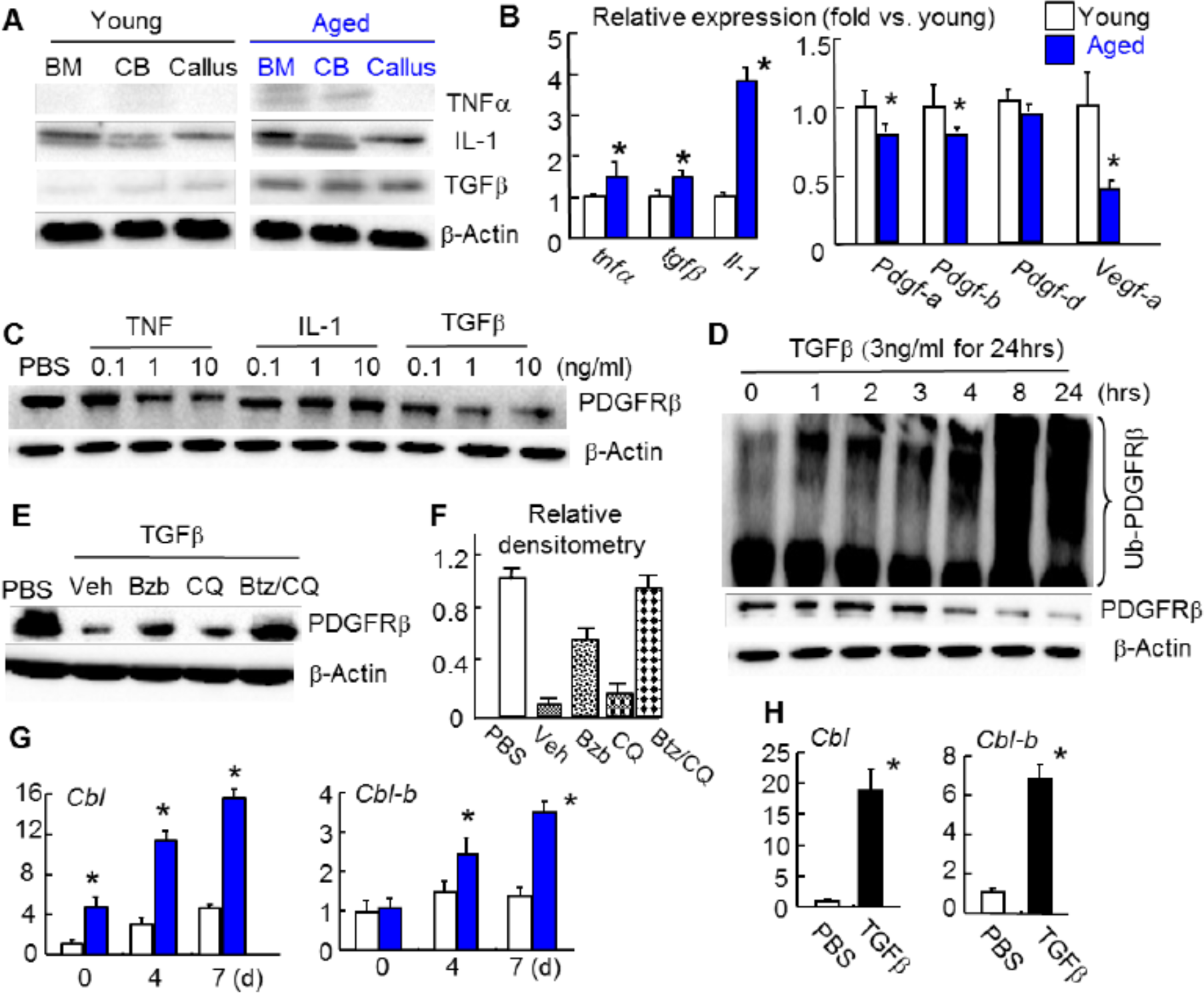
TGFβ induced PDGFRβ ubiquitination and proteasome degradation in aged mice. (A) Cytokines expressions in various tissues by WB. (B) Expressions of cytolines and PDGF family members in callus tissues by qPCR. (C-F) 3^rd^ passage of callus cells were used. (C) Cells were treated with TNF, IL-1 or TGFβ for 24 hours. PDGFRβ levels were measured by WB. (D) Cells were treated with TGFβ. Ub-PDGFRβ by Ub assays and total PDGFRβ levels were measured by WB. (E) Cells were treated with 3ng/ml TGFβ±3nM Btz, 10ug/ml CQ, and Btz/CQ for 24 hours. PDGFRβ level was measured by WB. (F) Densitometry of WB bands of (E). Fold changes over PBS. N=3. Values are mean±SD. (G) Expressions of Cbls in callus tissue by qPCR. N=3/group. Values are mean±SD. (H) Callus cells were treated 3ng/ml TGFβ for 3 days. Expressions of Cbls were measured by qPCR.

### Senescent cell derived TGFβ1 inhibit CaMPC growth through PDGFRβ

Since we investigated the molecular mechanism that TGFβ as the major SASP induced PDGFRβ proteasomal degradation, we further examined the biological function of senescent cell derived TGFβ on regulating CaMPC growth and PDGFRβ degradation. Data showed that TGFβ neutralizing Ab blocked the effect of aged CM on CaMPC growth, Ub-protein and PDGFRβ expression, which were inhibited by the PDGFRβ inhibitor, su16f (Fig. 3A&B). Data suggest that SCs in aged mice impaired fracture repair through regulating PDGFRβ via TGFβ.

**Fig.3.**
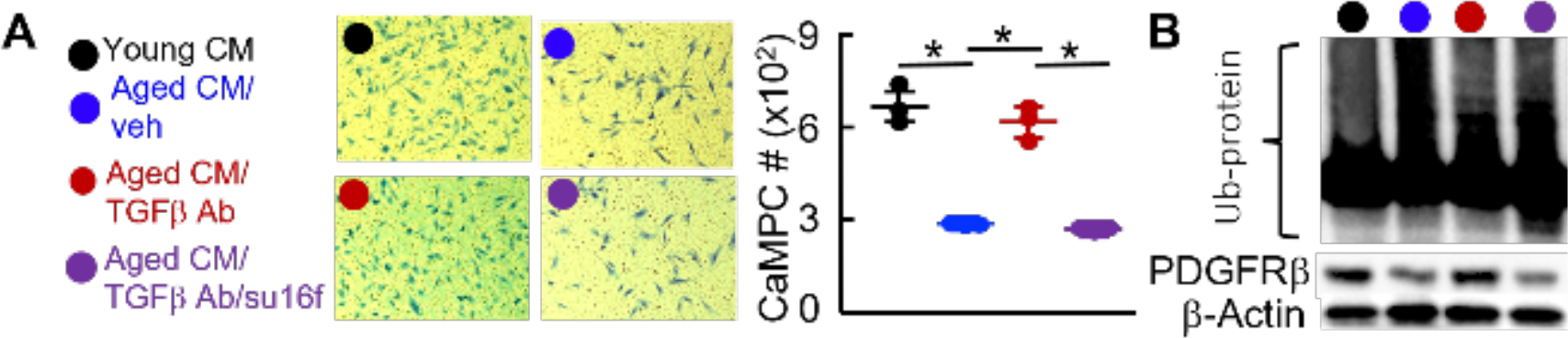
SCs derived TGFβ1 induces ubiquitination and proteasome degradation of PDGFRβ in aged mice. Fractured 3- and 20-m-old mice were sacrificed at d10. (A) CaMPCs were treated with young or aged callus CM ± Btz or CQ for 1 day. Ub-PDGFRβ was determined by an Ub assay. PDGFRβ protein levels were determined by WB. (B) PDGFRβ expression in TGFβ1 (3ng/ml)-treated CaMPCs ±Btz or CQ by WB.

## Discussion

SCs accumulate in multiple organs during aging^22^. Removal of SCs represents a promising therapeutic approach for age-related disorders^8^. Although SCs removal was reported to alleviate the aging phenotype in bone by regulating osteoblasts and osteoclasts through Sost and RANKL respectively^7^, the precise molecular mechanism has yet to be thoroughly examined. In the current study, we demonstrated that SCs increased protein ubiquitination and degradation in CaMPCs. Both senolytic drugs or Btz alone enhanced CaMPC growth and fracture repair in aged mice, but no addictive effect was observed when senolytic drugs and Btz were administered together, which infer that the effect of senescent cells on CaMPCs or fracture repair relies on UPS activity or adversely. Since proteasome inhibitor was reported to induce cellular senescence^18^ and we also found that Btz has no direct effect on cell senescence showed by expressions of senescent cell marker genes^6,23^, p16 and p21 by qPCR. Collectively, our data strongly imply that SCs impair fracture repair in aged mice by increasing UPS activity in CaMPCs.

Although numerous studies^6-8^ showed that senescent cells have detrimental effect on bone cells in aging related disease, including osteoporosis, osteoarthritis and aging fracture, there is still a big gap on the study of molecular mechanisms by which senescent cells regulating the function of neighboring bone cells through SASPs, which is a key feature of senescent cells. SASP is a complex mixture of tons of cytokines, chemokines, growth factor and matrix proteins, which means multiple key players are working together to affect the bone phenotype in aging related disease^4^ and also results in a big difficulty to investigate the exact molecular mechanism involved. We reported that proteasome inhibitor enhanced fracture repair in aged mice through inhibiting positive regulators for osteogenesis^13^. Senescent cells are the major sources chronic inflammatory cytokines in aged bones^9^, whose expressions were further boosted under acute injury^6^. Protein ubiquitination was closely associated with high expressions of inflammatory cytokines^9^, because most of E3 ligases are regulated by inflammatory response signal pathways, such as NF-κB pathway^14,24,25^. Through unbiased Ub-proteomics analysis, we found that PDGFRβ, which regulated both osteogenesis and angiogenesis was one of most ubiquinated proteins in MPCs (another manuscript to Biorxiv). Meanwhile, by searching the SASPs from aged mice, TGFβ was the most highly expressed one^6^, which has the higher possibility to decide the effects of senescent cells on neighboring cells. Collectively, senescent cell derived TGFβ regulate the ubiquitination of PDGFRβ in CaMPCs and contribute to the impaired fracture repair in aged mice. However, we will not exclude the possibility of SASPs, other than TGFβ contribute to the protein ubiquitination, because other increased inflammatory cytokines in callus of aged mice, such as TNFα or IL-1 can activate NF-kB signaling^26,27^, which in turn promote the expression of numerous E3 ligases^14,24,25^. Similarly, UPS has multiple targets in MPCs, because we also found other interesting proteins highly ubiquinated, such as Interferon-induced transmembrane protein (another manuscript to Biorxiv).

Overall, our study has revealed a new molecular mechanism whereby SCs regulate the function of CaMPCs during fracture repair in aged mice and opens avenues for optimizing treatment strategies with senolytic drugs for aging related diseases, such as aging fracture, by combination with the ligand of PDGFRβ, PDGF-BB.

## Materials and methods

### Animals

Young (4-month-old, 26-year-old in human) and aged (21-month-old, 62-year-old in human) C57BL/6J (WT) mice from National Institution of Aging were used. Mice were housed in micro-isolator technique rodent rooms. All animal procedures were approved by the University Committee on Animal Research at the University of Rochester.

### Tibial fracture procedure and animal treatments

Open tibial fracture procedures were performed according to standard operation procedure established in Center for Musculoskeletal Research^6^. In brief, an incision of 5 mm in length was made in the skin over the anterior side of the tibia after anesthesia. A sterile 27 G x 1.25-inch needle was inserted into the bone marrow cavity of the tibia from the proximal end, temporarily withdrawn to facilitate transection of the tibia using a scalpel at midshaft, and then reinserted to stabilize the fracture. The incision was closed with 5-0 nylon sutures. Fractures were confirmed by radiograph. Callus tissues were harvested on day 10, the time when soft callus is formed, following fracture procedure for cell preparation. Mice received slow-released extended release (XR) buprenorphine, 0.5 mg/kg, to control pain. Fractured mice were given D (Dasatinib, 5 mg/kg, Millipore Sigma, catalog CDS023389 by gavage) +Q (quercetin, 50 mg/kg, Millipore Sigma, catalog PHR1488 by gavage), Btz (Bortezomib, 0.6mg/kg by i.p.), su16f (PDGFRβ inhibitor 10mg/kg by gavage), TGFβ neutralizing Antibody 1D11 (2 μg/10 μL, R&D, catalog/clone MAB1835/1D11 by local injection) or vehicle at 1, 3, 5, and 7 dpf.

### Preparation of CaMPCs

Callus derived mesenchymal progenitor cells (CaMPCs) are obtained as described previously^6^. Briefly, callus tissues were dissected from the tibia, cut into small pieces (<1 mm3), and digested in 1 ml of Accumax solution (STEMCELL, 1 hour, room temperature). Cells were passed through a 35 um-filter and red blood cells were lysed with ammonium chloride (5 minutes, room temperature). Cells were cultured in basal medium (alpha-MEM medium containing 15% FBS). Cells that migrated from callus pieces were cultured in the basal medium to confluence, and CaMPCs in the 3rd-5th passage were used for experiments.

### Generation of conditioned medium and co-culture

For generating callus conditioned medium (CM)^6^, calluses harvested at 10 dpf were cut into pieces and cultured in aMEM (Minimum Essential Medium Eagle - Alpha Modification) medium containing 15% FBS (fetal bovine serum) for 2 day and CM is collected. For the co-culture experiments, CaMPCs growing on plastic well or 3D decellularized Wharton jelly matrix from the human umbilical cord (gifts from co-author, Brea Lipe’s lab)^16^ were treated with 30% CM ± 5uM su16f ± 3nM Btz ± 10ug/ml CQ ± 10ng/ml TGFβ neutralizing antibody (R&D) or PBS for 2 d and then subjected to the following assays: 1) Growth was determined by methylene blue staining in which the cells were fixed in 10% formalin and stained with 1% methylene blue solution. The percentage of positively stained area per well was calculated with Image J software on scanned images. 2) Cells were collected and subject to qPCR or western blots.

### Biomechanical testing

Fresh tibiae at 28 dpf were stored at –80°C after removing the stabilizing pins and soft tisses. The tibial ends were embedded in polymethylmethacrylate and placed on an EnduraTec system (Bose). A rotation rate of 10/s was used to twist the samples to failure or up to 80°. The bone strength parameter, maximum torque was analyzed following CMSR SOPs^6^.

### Quantitative real time qPCR

Callus tissues or cultured CaMPCs were subjected to RNA extraction with TRIzol, and cDNA was synthesized using the iSCRIPT cDNA Synthesis kit (BioRad). qPCR was performed with iQ SYBR Green Supermix using an iCycler PCR machine (BioRad). The fold change of gene expression was first normalized to actin and then normalized to the values in control group.

### Ubiquitination assay and western blot analysis

For ubiquitination assay^24^, CaMPCs were treated with the proteasome inhibitor MG 132 (10 μM) for 4 hours. Whole-cell lysates (200 μg protein/sample) were incubated with UbiQapture-Q Matrix (Biomol) by gentle agitation at 4°C overnight to pull down all ubiquitinated proteins according to the manufacturer’s instructions. After washing three times, captured proteins were eluted with 2× SDS-PAGE loading buffer and analyzed by Western blotting using anti-PDGFRβ antibody, as described previously (ref). For western blot analysis, callus tissues homogenized under liquid nitrogen or CaMPCs were extracted proteins with RIPA lysis buffer. Proteins were quantitated using a kit from Bio-Rad and loaded onto 10% SDS-PAGE gels and blotted with anti–TGFβ, TNFa, IL-1, PDGFRβ or actin antibodies. Bands were visualized using ECL chemiluminescence (Bio-Rad, catalog 1705061).

### Statistical analysis

Statistical analysis was performed using GraphPad Prism 5 software (GraphPad Software Inc., San Diego, CA, USA). Comparisons between two groups were analyzed using a 2-tailed unpaired Student’s t-test. One-way ANOVA and Tukey post-hoc multiple comparisons were used for comparisons among three or more groups.

